# Redox-dependent dimerization of PolDIP2 and a conserved ApaG-domain motif required for CHCHD2 interaction

**DOI:** 10.64898/2026.03.14.711777

**Authors:** Tran Vinh Hong Nguyen, Andreas Berner, Kazutoshi Kasho, Anais Lamy, Kieran Deane-Alder, Koit Aasumets, Namrata Chaudhari, Cuncun Qiao, Louise Leite Fernandes, Ronnie Berntsson, Sjoerd Wanrooij

## Abstract

PolDIP2 is a multifunctional mitochondrial protein implicated in redox regulation, mitochondrial proteostasis, and diverse mtDNA-associated processes, yet the principles underlying its regulation remain unclear. Crystallographic analysis revealed that PolDIP2 forms a redox-dependent disulfide-linked homodimer via a conserved Cys143 residue within its N-terminal YccV-like domain, and cellular and *in vitro* assays confirmed that this residue is essential for dimer formation. Oxidative stress enhanced dimerization of endogenous and ectopically expressed PolDIP2, and dimers were detected exclusively within mitochondria, requiring proper mitochondrial import. WT and C143A PolDIP2 overexpression produced similarly modest effects on mtDNA replication in cells, suggesting that dimerization has limited impact on mtDNA-associated processes. Proteomic analysis and biochemical validation identified both previously known and not yet characterized mitochondrial interactors of PolDIP2, and highlighted CHCHD2 as a specific binding partner. A conserved glycine-rich motif in the C-terminal ApaG/DUF525-like domain proved essential for this interaction, and disruption of the motif enhanced Cys143-dependent dimerization while abolishing CHCHD2 association, which preferentially occurs with monomeric PolDIP2. These findings define redox-controlled dimerization and a conserved ApaG-domain motif as key structural features shaping PolDIP2’s interaction state within mitochondria and provide a basis for exploring its roles in redox-sensitive mitochondrial pathways.

## INTRODUCTION

Mitochondria are central regulators of energy production and redox homeostasis. Oxidative phosphorylation generates reactive oxygen species (ROS), as a by-product, producing a redox-active environment that fluctuates with metabolic demand and stress. Proteins that function within mitochondria must therefore adapt to changes in oxidative conditions, yet the mechanisms that regulate the redox responsiveness, oligomeric state, or interaction networks of many mitochondrial proteins remain poorly understood^1,2^.

Polymerase δ-interacting protein 2 (PolDIP2/PDIP38) is suggested to be a multifunctional protein that localizes to both the nucleus and mitochondria^3^. Initially identified through interactions with DNA polymerase δ and replication factors^4^, PolDIP2 has since been implicated in processes including translesion DNA synthesis^5^, mtDNA maintenance^6^, oxidative signaling^7^, and metabolic regulation^6,8^. A substantial fraction of PolDIP2 resides in mitochondria, where it associates with mtDNA nucleoids^9^ and has been shown *in vitro* to enhance the polymerase activity of the primase–polymerase PrimPol^6^. Structurally, PolDIP2 contains an N-terminal mitochondrial targeting sequence, a YccV-like domain, and a C-terminal ApaG/DUF525-like domain^10^. These domains occur in protein families with roles in redox regulation and proteostasis, suggesting potential functional parallels^11^. Recent structural work showed the ApaG/DUF525 domain contains a conserved hydrophobic groove characteristic of substrate-binding pockets, supporting a role for PolDIP2 in mediating specific protein interactions within mitochondria^12^. Despite its broad interaction profile and diverse cellular roles, the molecular principles that govern PolDIP2’s regulation within mitochondria remain unclear.

In particular, it is unresolved whether PolDIP2 undergoes redox-dependent conformational changes or oligomerization, and how such features might influence its mitochondrial distribution or protein–protein interactions. The identity and specificity of PolDIP2’s mitochondrial binding partners also remain incompletely characterized. Among the regulatory proteins operating within the redox-active mitochondrial environment is CHCHD2, which maintains cristae organization^13^ and contributes to oxidative-stress responses^14^. CHCHD2 thus exemplifies the type of redox-sensitive pathway that PolDIP2 could interface with within mitochondria, although a direct involvement has not been established.

To address these gaps, we examined the biochemical and cellular properties of human PolDIP2, with the focus on how its structural domains contribute to redox responsiveness, oligomeric state, mitochondrial localization, and interaction specificity. We show that PolDIP2 forms a Cys143-dependent disulfide-linked dimer, that this dimerization is selectively enhanced under oxidative stress, and that dimer formation occurs exclusively within mitochondria. Recombinant monomeric and dimeric PolDIP2 stimulated PrimPol similarly, and WT and Cys143A overexpression caused comparable modest changes in mtDNA replication, indicating little influence of dimerization on mtDNA maintenance. Using proteomic and biochemical approaches, we identify CHCHD2 as a previously uncharacterized PolDIP2 interactor and demonstrate that a conserved glycine-rich motif within the C-terminal domain of PolDIP2 is required for this association. Together, these findings define key structural and redox-sensitive features that regulate PolDIP2 within mitochondria and reveal a molecular connection between PolDIP2 and the mitochondrial stress-responsive protein CHCHD2.

## MATERIAL AND METHODS

### Cloning of expression constructs

The full-length cDNA of PolDIP2 WT and its variants with a Flag tag was cloned into pcDNA5/FRT/TO vector (Invitrogen), as previously described^15^. The mutant constructs were generated by site-directed mutagenesis using PfuUltra high-fidelity (Agilent) according to the manufacturer’s instruction, and all resulting plasmids were validated by Sanger sequencing.

### Oligonucleotides

For primer extension assays a 5′-TET fluorophore-labelled 25-mer oligonucleotide primer (5′-TET-ATAGGGGTATGCCTACTTCCAACTC-3′) was annealed with a 70-mer oligonucleotide template (5′-GAGGGGTATGTGA TGGGAGGGCTAGGATATGAGGTGAGTTGAGTGGAGTTGGAAGTAGGCATACCCT-3′) at 1:1.2 primer: template ratio in the presence of 100 mM NaCl by incubating at 95 °C for 5 min and cooling down to room temperature. The annealed primer-template was kept at -20 °C for short term storage.

### Cell culture and compound preparation

Inducible cell lines expressing PolDIP2 variants were established by using Flp-In™ T-REx™ HEK293 host cell-line (Thermo Fisher Scientific) as described essentially in the previous work^15^. Flp-In T-REx 293 cells were maintained in DMEM high glucose + GlutaMAX (Life Technologies) supplemented with 10 % Fetal Bovine Serum (Sigma-Aldrich), 1 mM sodium pyruvate, 50 μg/ml uridine and Pen/Strep at 37 °C in a humidified incubator with 7% CO_2_ atmosphere. The expression of transgene was induced by adding doxycycline (Dox) to the growth medium at the final concentration of 10 ng/ml. At this concentration, Dox gives a stable long-term expression of the transgene and is nontoxic for mitochondrial functions.

Parental and PolDIP2 K.O HAP1 cells were purchased from Horizondiscovery.

2’,3’-dideoxycytidine (ddC, Abcam) was freshly prepared in distilled water prior to cell treatment. Rotenone (Sigma-Aldrich) was dissolved in DMSO and kept at -20°C in small aliquots. Mitochondrial uncoupler BAM15 (Sigma-Aldrich) and cycloheximide (CHX, Sigma-Aldrich) were dissolved in distilled water and kept at -20°C in small aliquots for short-term storage. Hydrogen peroxide (H_2_O_2_, Sigma-Aldrich) was aliquoted in small volume and stored at 4 °C.

### siRNA silencing

Cells were reverse transfected with siRNA upon seeding onto 6-well plates. Briefly, a mixture of Lipofectamine® RNAiMAX (Invitrogen) and 90 pmol PolDIP2 siRNA ((#4392420, Thermo Fisher) or scrambled siRNA (#4390844, Thermo Fisher, as negative control) was prepared in OptiMEM® medium (Gibco) according to manufacture and added dropwise to the cells. 72 hours after transfection, cells were harvested for protein extraction and analysed by immunoblotting.

### Protein extraction and immunoblotting

Cell cultured medium was aspirated and cells were then collected in ice-cold 1X PBS before being lysed in precooled NP-40 lysis buffer (10 mM Tris/Cl pH 7.5, 150 mM NaCl, 0.5 mM EDTA, 0.5 % Nonidet™ P40 Substitute, and supplemented with 1x Halt protease inhibitor cocktail (ThermoFisher Scientific)) for 20 min on ice. The supernatant containing total protein extract was collected after maximum-speed centrifugation for 15 min at 4 °C. The protein samples were quantified using Pierce BCA assay kit (Thermo Scientific). 10-20µg of total protein extract was dissolved in 1x Laemmli Sample Buffer (Biorad) with β-mercaptoethanol and boiled at 95 °C for 5 min prior to being separated on 4-20 % Mini-PROTEAN® pre-casted gel (Bio-Rad) at 100V and transferred to 0.45 μm nitrocellulose membranes (GE Healthcare Life Sciences) using a Mini-Protean electrophoresis system (Bio-Rad) for 1 h at 4°C. For non-reducing SDS-PAGE, protein extract was dissolved in 1x Laemmli Sample Buffer (Biorad) and kept at RT before use.

The membranes were then blocked in 5 % non-fat milk for 1h at RT prior to incubation with primary antibody overnight at 4 °C, followed by incubation with horseradish peroxidase-conjugated-secondary antibodies for 1h at RT. All incubation and washes were performed in Tris-buffered saline with 0.1% Tween-20. The antibodies used and their dilutions are listed in Table 1. Chemiluminescent detection was performed using Super Signal™ West Pico Chemiluminescent Substrate (Thermo Scientific) on ChemiDoc Touch Imaging System (Bio-Rad).

**Table 1.** Antibodies used in the study.

| **Primary Antibody** | **Host** | **Cat.number** | **Dilution in WB** | **Purchased** |
| --- | --- | --- | --- | --- |
| Flag M2 | Mouse | F1804 | 1:2000 | Sigma Aldrich |
| PolDIP2 | Rabbit | Ab181841 | 1:2000 | Abcam |
| Beta-actin | Mouse | 66009-1-Ig | 1:10000 | Proteintech |
| GRPEL2 | Rabbit | NBP1-85099 | 1:5000 | Novus Biologicals |
| Lamin A/C | Rabbit | 10298-1-AP | 1:5000 | Proteintech |
| GAPDH | Rabbit | MA5-15738 | 1:5000 | Invitrogen |
| TOM-20 | Rabbit | 11802-1-AP | 1:5000 | Proteintech |
| CLPX | Rabbit | PA5-79052 | 1:5000 | Invitrogen |
| CHCHD2 | Rabbit | 19424-1-AP | 1:2000 | Proteintech |
| HSP60 | chicken | PA5-143571 | 1:5000 | Invitrogen |
| LONP1 | Rabbit | 15440-1-AP | 1:5000 | Proteintech |
| **Secondary Antibody** | **Host** | **Cat.number** | **Dilution in WB** | **Purchased** |
| Goat-anti mouse IgG HRP | Goat | #31430 | 1:10000 | Thermo Scientific |
| Goat-anti rabbit IgG HRP | Goat | #31460 | 1:10000 | Thermo Scientific |
| Goat-anti chicken IgG HRP | Goat | # A16054 | 1:10000 | Thermo Scientific |

### Co-Immunoprecipitation

Cells were seeded on 10-cm plates and reached approximately 90% confluency at the day of harvesting. After 24 hours of induction with 10 ng/mL doxycycline, cells were collected in ice-cold 1X PBS before being lysed in precooled NP-40 lysis buffer (10 mM Tris/Cl pH 7.5, 150 mM NaCl, 0.5 mM EDTA, 0.5 % Nonidet™ P40 Substitute, and supplemented with 1x Halt protease inhibitor cocktail (ThermoFisher Scientific)) for 30 min on ice. The supernatant containing total protein extract was collected after maximum-speed centrifugation for 15 min at 4°C. Exogenous Flag-tagged PolDIP2 was enriched using ChromoTek DYKDDDDK Fab-Trap® Agarose beads (Proteintech) as described in the manufacture’s instruction. Briefly, cell extract was added to the beads, and the mixture were rotated end-over-end for 1 hour at +4°C. The beads were then sedimented by centrifugation at 2,500x g for 5 min at +4°C and washed several times with Wash buffer (10 mM Tris/Cl pH 7.5, 150 mM NaCl, 0.5 mM EDTA, **0.05%** Nonidet™ P40 Substitute). The bound proteins were subsequently eluted by pipetting the beads up and down several times in 200 mM glycine pH 2.5, followed by immediate neutralization with 1 M Tris pH 9.0. The eluate was then analysed on SDS-PAGE and Western Blotting as described above or was subjected to Mass photometry analysis.

### Cell fractionation

To determine the sub-cellular localization of PolDIP2, cells were grown 10-cm plates to 70% confluence prior to transfection with different PolDIP2 variants for at least 24 hours. Reverse and forward transfection using Lipofectamine 2000 (Invitrogen) were performed according to manufacturer’s instructions in HAP1 and Flp-In™ T-REx™ HEK293, accordingly. Cells were harvested, washed twice with ice-cold 1xPBS, and then resuspended in 0.1x homogenization buffer (HG) (4 mM Tris-HCl pH 7.8, 2.5 mM NaCl, 0.5 mM MgCl_2_) for 10 min on ice. Cells were then homogenized in a 1mL glass Wheaton tight-fitting dounce (Active Motif) and sub-cellular fractions (nuclear, cytosolic, and mitochondrial fractions) were obtained after differential centrifugation. Proteins from these different fractions were extracted by incubating in precooled NP-40 lysis buffer for 20 min on ice and collecting the supernatant after maximum-speed centrifugation for 15 min at 4°C.

### Protein purification

PrimPol was purified as described previously^16,17^. Human truncated PolDIP2 (AA 59-355) was cloned into pET28a(+) vector in frame with the C-terminal TEV cleavable 6xHis tag. Plasmid was transformed into *E. coli* Rosetta DE3 chemical competent cells. An overnight culture was grown and used to inoculate 1.5 L of TB without antibiotics. Culture was grown at 30°C in a Lex bioreactor system until the OD_600_ reached 0,6. Temperature was reduced to 20 °C and expression was induced with 1 mM IPTG. After 20h incubation cells were collected by centrifugation at 4000 × g for 15 min at 4 °C. Pellets were resuspended in 30 ml lysis buffer (50 mM Tris-HCl pH 8.0, 300 mM NaCl, 20 mM Imidazole, 0,5% Tween20, 1 µM Pepstatin, 0.1 mM AEBSF, 1 mM E64, 1 mM DTT and DNAse). Cells were lysed by sonication, and the supernatant was cleared by centrifugation at 25.000 x g at 4 °C for 1h. The supernatant was filtered through a 0.45 µm filter and applied to 2 ml pre-equilibrated (Buffer 1: 50 mM Tris-HCl pH 8.0, 300 mM NaCl, 20 mM Imidazole) Ni-NTA resin. The column was washed with 20 ml buffer 1, followed by 10 ml buffer 2 (buffer 1 with 1 M NaCl) and 10 ml buffer 3 (buffer 1 with 40 mM Imidazole). Protein was eluted with 10 ml elution buffer (buffer 1 with 330 mM Imidazole). 250 µL TEV protease (2mg/ml) was added and incubated for 1h at RT to cleave the 6xHis tag. Protein was dialysed against buffer 4 (50 mM Tris-HCl pH 8.0, 100 mM NaCl) over night at 4 °C. After dialysis, protein was loaded on 2ml pre-equilibrated Ni-NTA column again and flow through was collected. Sample was concentrated to 1.5 ml with ∼11 mg/ml protein. For dimerization, 5 mM K_4_[Fe(CN)_6_] was added and incubated over night at RT. The next day, the protein sample was filtered using a 0.22 µm filter and injected onto on a Superdex 200 Increase 10/300 GL equilibrated in buffer 4. Monomers and dimers were separated and analysed on SDS-PAGE. Fractions corresponding to monomer/dimer were aliquoted, frozen in liquid nitrogen and stored at 80 °C.

### Crystallization and structure determination

Protein crystals were obtained with the sitting drop vapor diffusion method at 20°C. PolDIP2 crystallized in a 1:1 ratio with 15% (w/v) PEG 3350, 0.1 M Hepes (pH 7.7) and 0.075 M MgCl_2_ with a concentration of PolDIP2 at 4 mg/ml. Crystal was cryo-protected with 25% ethylene glycol before flash-freezing in liquid nitrogen. X-ray diffraction data was collected on BioMax, MaxIV, Sweden. The data was processed using XDS ^18,19^.

The crystal had the space group *P*2_1_2_1_2_1_ and contained 2 copies of the protein in the asymmetric unit. The phase-problem was solved using molecular replacement with the previously published structure of PolDIP2 (PDB code: 6Z9C) in PHENIX phaser ^20,21^. The structures were built in Coot ^22^ and refined at 2.3 Å in PHENIX refine ^23^, to R_work_/R_free_ values of 25.3/30.7%. Further refinement statistics can be found in Table S2.

### Primer Extension Assay

PrimPol primer extension experiments were performed as previously described^6^. Briefly, reactions containing 15 nM of hybridized 5′-TET-perimer/template and indicated concentrations of PrimPol and PolDIP2 monomer/dimer were prepared on ice and transferred to 37 °C for 20 min. The reactions were stopped by addition of stop solution. DNA products of primer extension assays were loaded on 10% polyacrylamide gels containing urea and imaged using a Typhoon 9400 scanner (Amersham bioscience). To check dimerization status of PolDIP2 after primer extension assays, the reactions were added 1x Laemmli Sample Buffer (Biorad) without β-mercaptoethanol and analysed on SDS-PAGE.

### Mass photometry

Protein samples were diluted in a dilution buffer (20mM Tris pH 8.0, 0.5 mM EDTA, 200mM NaCl, 2mM DTT) to a working concentration between 10 to 100 nM, depending on the dissociation properties of individual sample.

Microscope coverslips (No. 1.5H, 24×50 mm) were sequentially rinsed with isopropanol and Milli-Q water three times, which followed by drying with a clean compressed-air stream. The CultureWellTM gaskets (GBL103250-10EA, Sigma-Aldrich) were cut to cover an area containing four wells (3 mm diameter, 1 mm deep) and placed in the centre of the clean coverslip, assuring a tight fit through gentle pressure. A droplet of immersion oil (Carl Zeiss™ Immersol™518F, Fisher scientific) was applied to the objective of the flow-chamber of OneMP Mass Photometer (Refeyn Ltd, Oxford, UK), and the prepared coverslip-gasket assembly was mounted and stabilized with small magnets. Each protein was measured in a new well. The fresh dilution buffer was loaded in wells of the gasket and focus was determined by the autofocus system. For each measurement, 15 µl of protein samples was loaded into a well. Data acquisition was performed using AcquireMP (Refeyn Ltd, v 2.4.1) and Movies lasting 60 seconds were recorded. All MP movies were processed and analysed using DiscoverMP (Refeyn Ltd, v 2.4.2). Bovine Serum Albumin was used as the standard sample.

### Two-dimensional agarose gel electrophoresis (2D-AGE) and Southern blotting

The 2D-AGE analysis of mitochondrial DNA replication intermediates was performed as described previously^15,24^. Briefly, 10 µg of total DNA was digested with 4 µl of FastDigest *Hinc*II (Thermo Fisher Scientific) for ∼16 h at 37°C and separated over a 0.4% agarose gel in 1 × TBE without EtBr at low voltage (20–30 V) for 16 hours. The gel was stained with 1 µg/ml EtBr, the lanes were cut out and a 0.95% agarose gel containing 0.5 µg/ml EtBr was cast around the gel slab. Second-dimension gel was run at 110 V in 1 × TBE buffer with 0.5 µg/ml EtBr at 4°C until the linear fragments were ∼1 cm before the end of the gel. Southern blotting was performed as described before and the membrane was hybridized with a radioactively labelled probe for nts 37–611 of human mtDNA. Radioactive signals were captured and quantified using phosphor screens and a phosphor imager (Amersham Typhoon).

## RESULTS

### PolDIP2 dimerization occurs *in vitro* and in cultured human cells

PolDIP2 has been proposed to be involved in several cellular pathways, including DNA replication, mitochondrial metabolism, and redox regulation. While many of these functions are supported by experimental data, they appear to be context-dependent, and a unifying molecular mechanism has yet to be established. To gain insight into its biochemical properties and regulatory potential, we analyzed PolDIP2 in both *in vitro* and cellular systems. During these studies, we unexpectedly observed higher-molecular-weight forms of PolDIP2, suggesting self-association. Since oligomerization can affect protein activity, stability, and localization, we sought to characterize the oligomeric state of PolDIP2 under physiological and experimental conditions.

To determine whether PolDIP2 forms oligomers in cells, we analyzed endogenous PolDIP2 in HEK293T cells by SDS-PAGE under non-reducing conditions (in the absence of β-mercaptoethanol). PolDIP2 contains an N-terminal mitochondrial targeting sequence (MTS) that is cleaved upon import, producing a mature ∼37 kDa form. Western blot analysis revealed the expected ∼37 kDa band and an additional, slower-migrating species, at ∼85 kDa (**Fig. 1A**). This higher-molecular-weight band disappeared under reducing conditions (in the presence of β-mercaptoethanol), suggesting it represents a disulfide-linked oligomer. Despite being less abundant than the monomer, the oligomer was similarly reduced upon RNAi knockdown, indicating that both species are specific forms of PolDIP2 (**Fig. S1A**).

**Figure 1.**
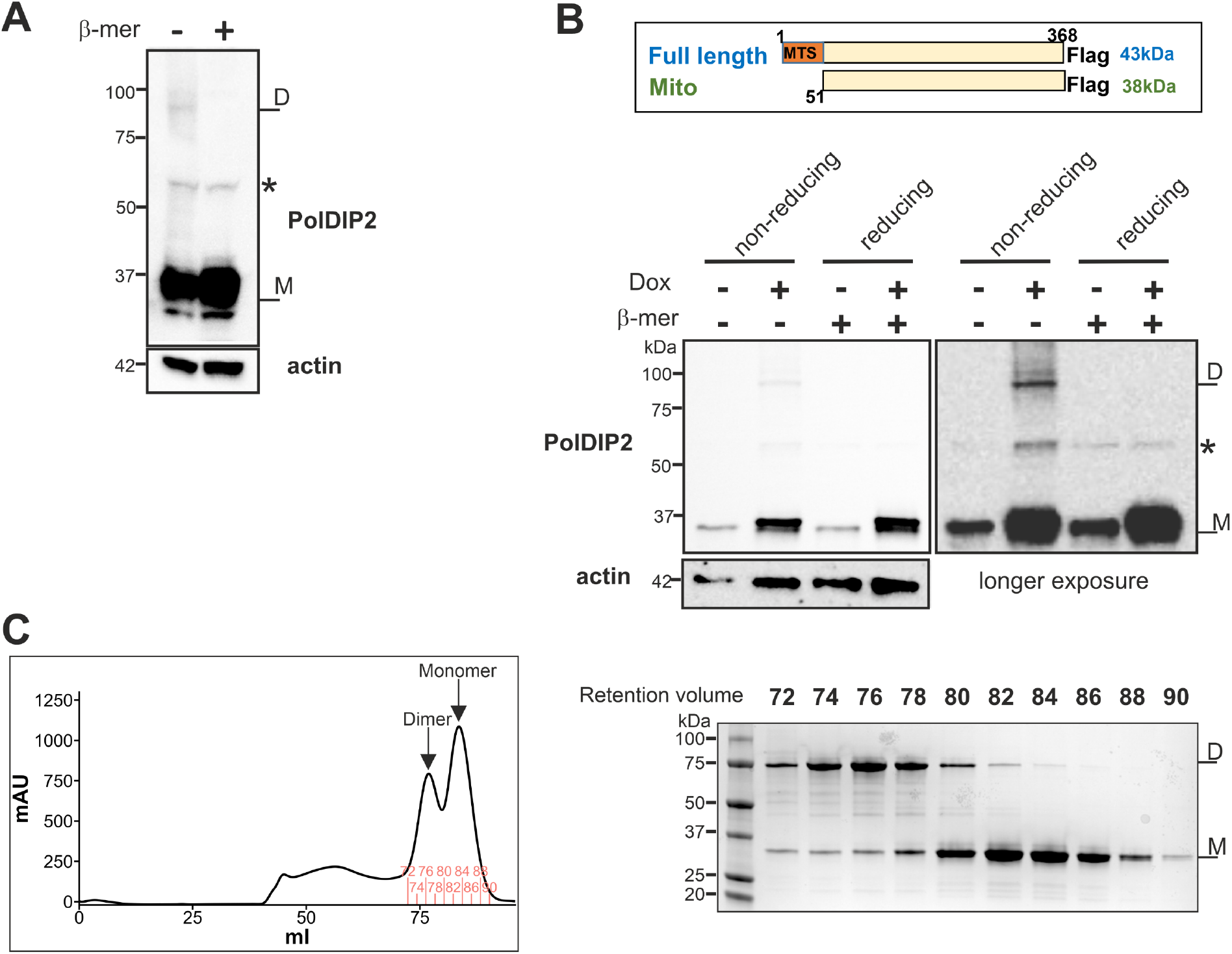
Human PolDIP2 dimerizes *in vitro* and in cultured human cells. **(A)** Western blot analysis using an anti-PolDIP2 antibody reveals two distinct bands, representing different endogenous PolDIP2 species: M as monomer, and D as dimer. The slower-migrating band is reduced upon treatment with β-mercaptoethanol (reducing), suggesting it corresponds to a disulfide-linked dimer. **(B)** Schematic representation of Flag-tagged PolDIP2 expressed in our cell model. Full-length PolDIP2 is processed upon import into mitochondria, where removal of its MTS yields the mature protein. Western blot analysis using an anti-Flag antibody detects C-terminally Flag-tagged PolDIP2 expressed in Flp-In™ 293 T-REx cells, with and without doxycycline induction, under reducing and non-reducing conditions. Both short (left) and long (right) exposures of the blot are shown. **(C)** Purified recombinant human PolDIP2 elutes as two distinct peaks during size-exclusion chromatography (left panel). Selected fractions were analyzed by SDS-PAGE (without β-mercaptoethanol in the loading buffer) and visualized with Instant Blue (right panel). Earlier fractions correspond to dimerized PolDIP2 (∼90 kDa), while the monomer eluted in the later fractions. *non-specific band.

To further investigate PolDIP2 oligomerization in living cells, we established a HEK293 FlpIn T-Rex cell line that expresses PolDIP2 with a C-terminal Flag tag upon doxycycline (Dox) addition. The overexpression of PolDIP2 was detected with both anti-Flag **(Fig. S1B)** and anti-PolDIP2 antibodies (**Fig. 1B**), confirming the conditional expression under doxycycline induction. Like endogenously expressed PolDIP2 (**Fig. 1A**), the PolDIP2-Flag migrated partly as higher-molecular-weight band that disappeared under reducing conditions (**Fig. 1B**).

To investigate the oligomeric state of human PolDIP2, we analyzed and purified recombinant truncated protein (residues 59-355, ∼34 kDa) expressed in *E. coli*. The recombinant PolDIP2 was incubated with potassium ferricyanide (K_3_Fe(CN)_6_) before being subjected to size exclusion chromatography (SEC) column (**Fig. 1C**). The majority of PolDIP2 eluted in later fractions (**Fig. 1C,** retention volume 80-90 ml) and migrated as a monomer on SDS-PAGE (∼34 kDa). However, on non-reducing SDS-PAGE, a portion of PolDIP2 appeared as an ∼75 kDa band, corresponding to earlier SEC fractions (**Fig. 1C,** retention volume 72-80 ml), consistent with an oligomeric form. This SEC-based separation enabled the enrichment of both monomeric and oligomeric PolDIP2 species. Our mass photometry results confirmed that the higher-molecular weight species correspond to PolDIP2 dimers (**Fig. S1C**). Similarly, overexpressed PolDIP2-Flag enriched from cultured cells also contained two distinct populations corresponding to monomeric and dimeric forms (**Fig. S1C**).

Together, these results demonstrate that PolDIP2 can form disulfide-linked dimers both in cells and *in vitro*, providing a foundation for exploring the functional relevance of its oligomerization.

### Cysteine 143 is important for PolDIP2 dimer formation via a disulfide bond

To investigate the molecular basis of PolDIP2 dimerization, we solved the crystal structure of purified human PolDIP2 (residues 59-355) to 2.3Å. The structure revealed that Cysteine 143, located within the conserved YccV domain, mediated dimerization through the formation of an intermolecular disulfide bond between two PolDIP2 monomers (**Fig. 2A**). Because sequence alignment also identified Cys266 as a highly conserved cysteine within the ApaG domain (**Fig. 2B**), we next examined the contribution of both residues to dimer formation in cells. To this end, we generated doxycycline-inducible HEK293 cell lines expressing PolDIP2 variants in which Cys143 or Cys266 was individually substituted with alanine.

**Figure 2.**
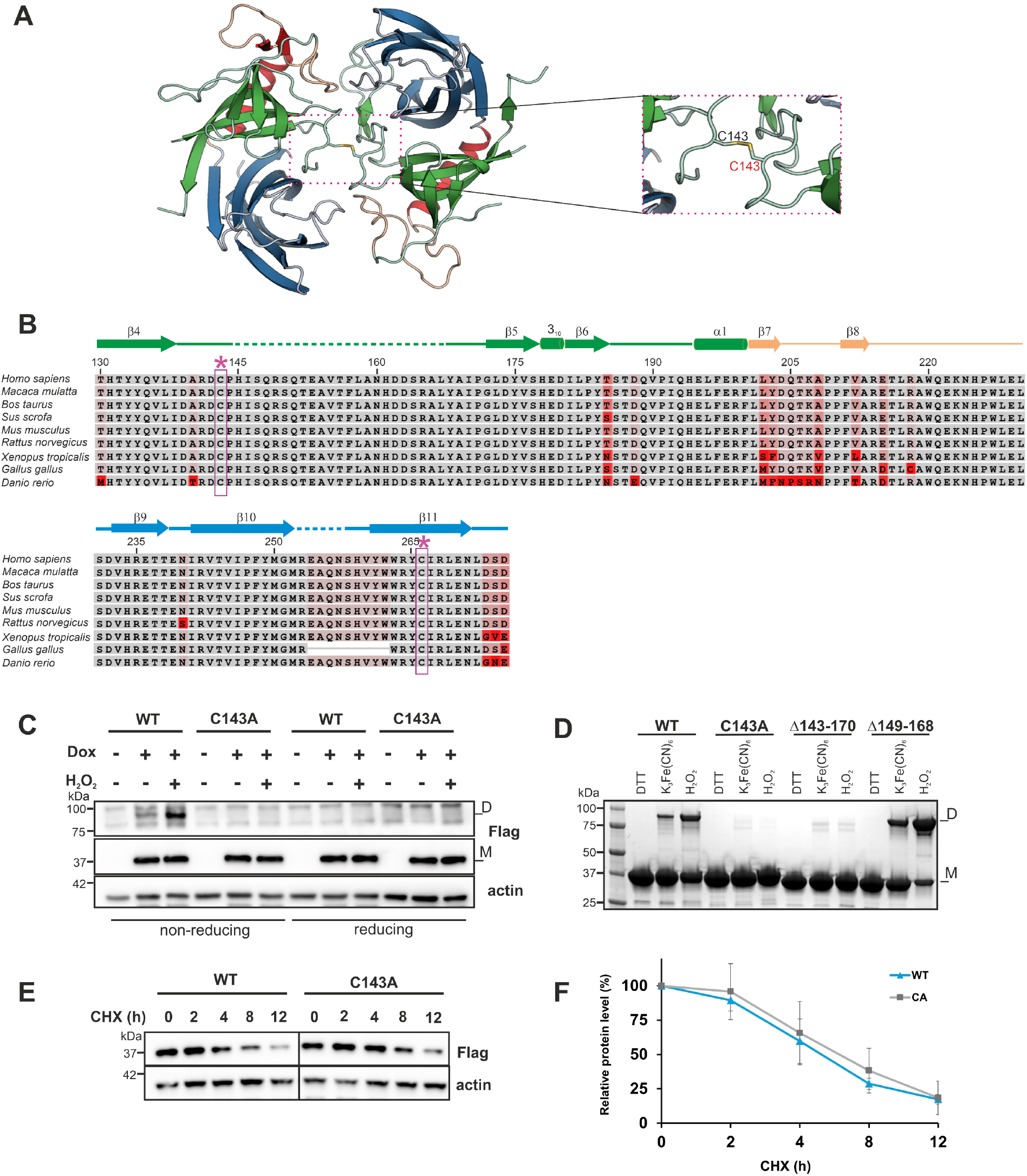
Cysteine 143 mediates disulfide bond-dependent dimerization of human PolDIP2 in cells and *in vitro*. **(A)** The crystal structure of dimeric human PolDIP2 reveals that Cys143 residues from each monomer form an intermolecular disulfide bond at the dimer interface. The left panel shows the overall structure of a dimer while the right panel is a close-up of the dimer interface. The structure shows symmetric pairing of the YccV domains and highlights the spatial proximity of Cys143 side chains, enabling the formation of covalent disulfide linkage. YccV-like domain is shown in green and ApaG-like domain is shown in blue, the two domains are connected by a linker region indicated in peach. **(B)** Protein sequence alignment of PolDIP2 from select vertebrate species. Two highly conserved cysteines, Cys143 in the YccV-like domain and Cys266 in the ApaG-like domain, are indicated with *. Conserved residues are shown in grey, and the shades of red colour indicate conservation levels of other residues across select species. Residue numbers correspond to human full-length PolDIP2 sequence. **(C)** Western blot analysis using anti-Flag antibody in Flp-In™ 293 T-REx cells expressing wild-type (WT) and C143A PolDIP2 variants. Only WT PolDIP2 shows dimer formation under non-reducing conditions and dimerization of the WT protein increases upon cell exposure with hydrogen peroxide. Cells were analyzed with or without doxycycline transgene induction (–/+), and actin served as a loading control. (**D**) SDS-PAGE of purified recombinant PolDIP2 variants (WT, C143A, Δ143–170, and Δ149–168) in reducing (DTT) or oxidative condition induced by potassium ferricyanide K_3_Fe(CN)_6_ or H_2_O_2_ visualized by Coomassie staining. (**E**) Representative Western Blots of PolDIP2 overexpressing cells treated with 10 µg/mL of cycloheximide (CHX). The protein levels of exogenous WT PolDIP2 and mutant were monitored over the time course upon CHX addition and detected with anti-Flag antibody. (**F**) Quantification of WT PolDIP2 and C143A mutant protein levels in (E) shows comparable turnover rates of the two PolDIP2 variants.

Consistent with observations for the wild-type protein, the C266A PolDIP2 mutant formed dimers in cells, as evidenced by the presence of a slower-migrating ∼85 kDa band on non-reducing SDS-PAGE (**Fig. S2A**). This band disappeared under reducing conditions, confirming the disulfide-linked nature of the dimer (**Fig. S2A**). On the contrary, the PolDIP2 C143A mutant variant was unable to form dimers and only the monomeric C143A PolDIP2 variant was visible in non-reducing conditions (**Fig. 2C**).

To confirm the role of Cysteine 143 *in vitro*, we analyzed recombinant PolDIP2 mutants (C143A, Δ143–170, and Δ149–168). Dimerization of purified monomeric wild-type PolDIP2 could be induced by oxidative conditions using potassium ferricyanide or H_2_O_2_ (**Fig. 2D**). In contrast, both C143A and Δ143–170 mutants failed to form dimers, while the Δ149–168 variant retained dimerization capacity, reinforcing the specific requirement of C143 for disulfide bond-mediated dimer formation. Together, these results establish Cysteine 143 as a critical determinant of PolDIP2 dimerization both in cells and *in vitro*.

As disulfide bonds can contribute significantly to protein stability, we asked whether disrupting the Cys143-dependent disulfide affects PolDIP2 turnover. Cells overexpressing WT PolDIP2 or the C143A mutant were treated with Cycloheximide, an inhibitor of protein synthesis, and protein levels were monitored over time (**Fig. 2E and F**). The C143A variant displayed a half-life comparable to WT (t_1/2_ ∼6 hours; **Fig. 2F**), indicating that loss of the disulfide bond does not appreciably alter steady-state stability. This suggested that Cys143 is specifically required for redox-responsive dimerization rather than for general folding or turnover of PolDIP2. These findings prompted us to test whether cellular oxidative stress modulates PolDIP2 dimer formation.

### ROS-induced oxidative stress enhances PolDIP2 dimerization

Given that endogenous PolDIP2 dimers were present at relatively low levels in HEK293T cells under basal conditions, we investigated whether their formation could be stimulated by specific cellular stressors. As demonstrated in **Figure 2A**, PolDIP2 dimerization was mediated by a disulfide bond between two cysteine residues, and this interaction can be induced in recombinant protein under oxidative conditions (**Fig. 2D**). To determine whether oxidative stress promotes dimerization in cells, we treated HEK293T cells with hydrogen peroxide (H_2_O_2_) and assessed endogenous PolDIP2 dimer levels. While baseline dimerization was already detectable, H_2_O_2_ treatment led to a concentration-dependent increase in PolDIP2 dimer formation (**Fig. 3A**). As a positive control, we monitored GRPEL2, a mitochondrial protein known to undergo redox-sensitive oligomerization and observed a similar pattern of H_2_O_2_-induced dimerization.

**Figure 3.**
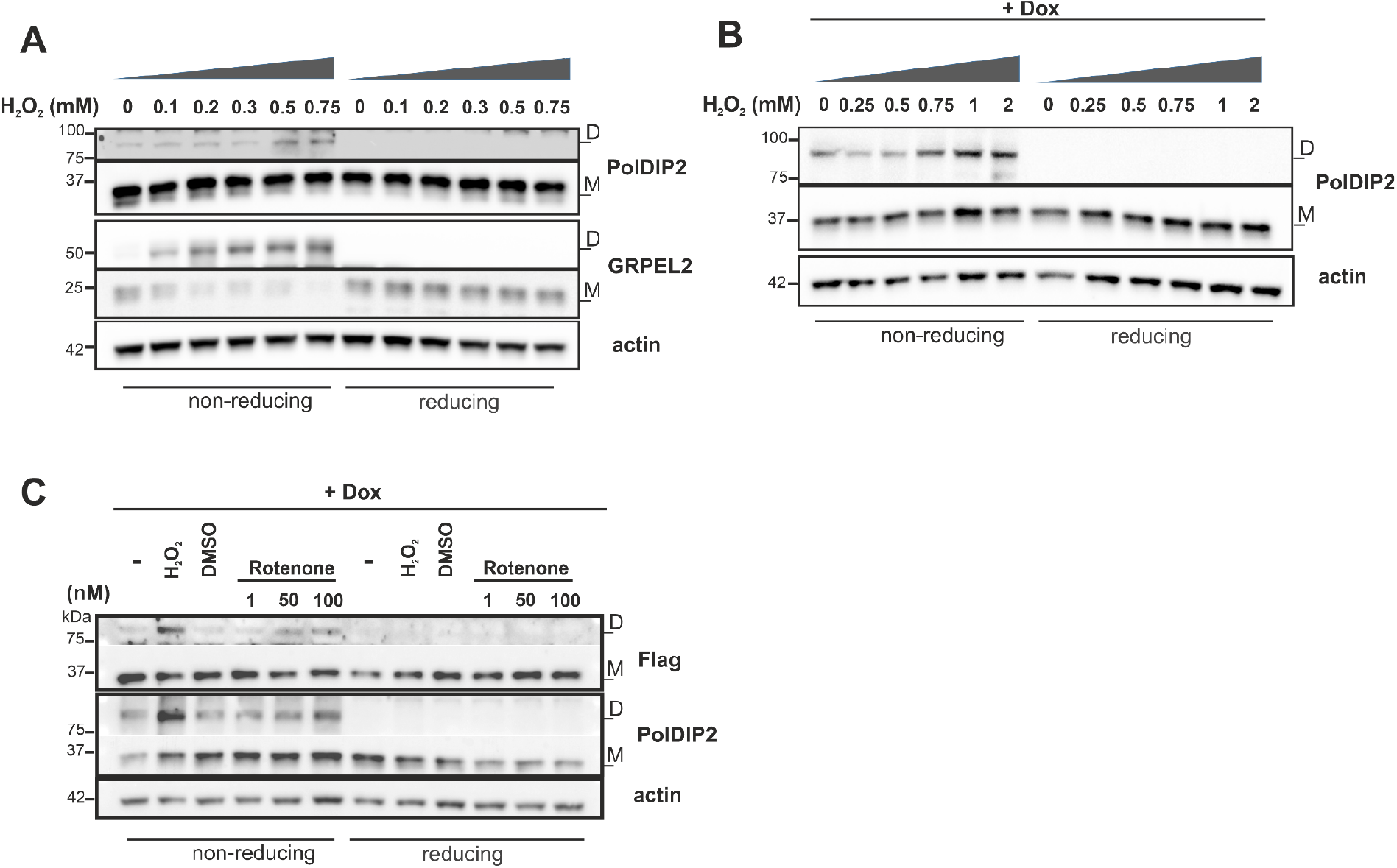
Oxidative stress enhances PolDIP2 dimerization in HEK293T cells. **(A).** Western blot analysis showed that H_2_O_2_ treatment induced dimer formation of endogenous PolDIP2. The gels were run under non-reducing and reducing conditions. GRPEL2, a redox-sensitive mitochondrial protein, was used as a positive control for H_2_O_2_-induced dimerization, and actin served as a loading control. **(B).** H_2_O_2_ treatment induced exogenous PolDIP2-Flag dimerization in doxycycline-inducible Flp-In 293 Trex cells. **(C).** Flp-In Trex PolDIP2 cells were treated with vehicle DMSO, H_2_O_2_ (15 min) or rotenone for 24 hours.

Consistent with the behavior of endogenous protein, exogenous Flag-tagged PolDIP2 also showed enhanced dimerization following H_2_O_2_ treatment (**Fig. 3B**). Furthermore, exposure to rotenone, an inhibitor of mitochondrial complex I that increases intracellular reactive oxygen species (ROS), also promoted PolDIP2 dimerization, albeit to a lesser extent than H_2_O_2_ (**Fig. 3C**). In contrast, treatment with 2’,3’-dideoxycytidine (ddC), a nucleoside analog that disrupts mitochondrial DNA replication, did not enhance PolDIP2 dimer formation (**Fig. S2B**). Importantly, H_2_O_2_-induced exogenous PolDIP2 dimerization was absent in cells expressing the Cys143A PolDIP2 variant, indicating the importance of Cys143 for disulfide bond formation of human PolDIP2 dimer under oxidative conditions (**Fig. 2C**). Taken together, these results indicate that PolDIP2 dimerization is selectively upregulated in response to oxidative stress mediated by elevated ROS levels.

### Cellular localization of PolDIP2 monomer and dimer

Although human PolDIP2 contains a canonical N-terminal mitochondrial targeting sequence, previous studies have reported its localization to various other subcellular compartments. While these observations suggest potential context-dependent distribution, the functional relevance of these alternative localizations remains to be fully clarified. Thus, we next investigated the subcellular localization of PolDIP2 dimerization using a cell fractionation assay in Flp-In 293 T-Rex cells. As shown in **Figure 4A**, while the monomeric form of PolDIP2 was detected in both the cytoplasmic and mitochondrial fractions, the dimeric form was exclusively localized to the mitochondria. This mitochondrial localization of the PolDIP2 dimer was further supported by the loss of the dimeric species upon treatment with the mitochondrial uncoupler BAM15 which disrupts protein import into mitochondria (**Fig. S3A**).

**Figure 4.**
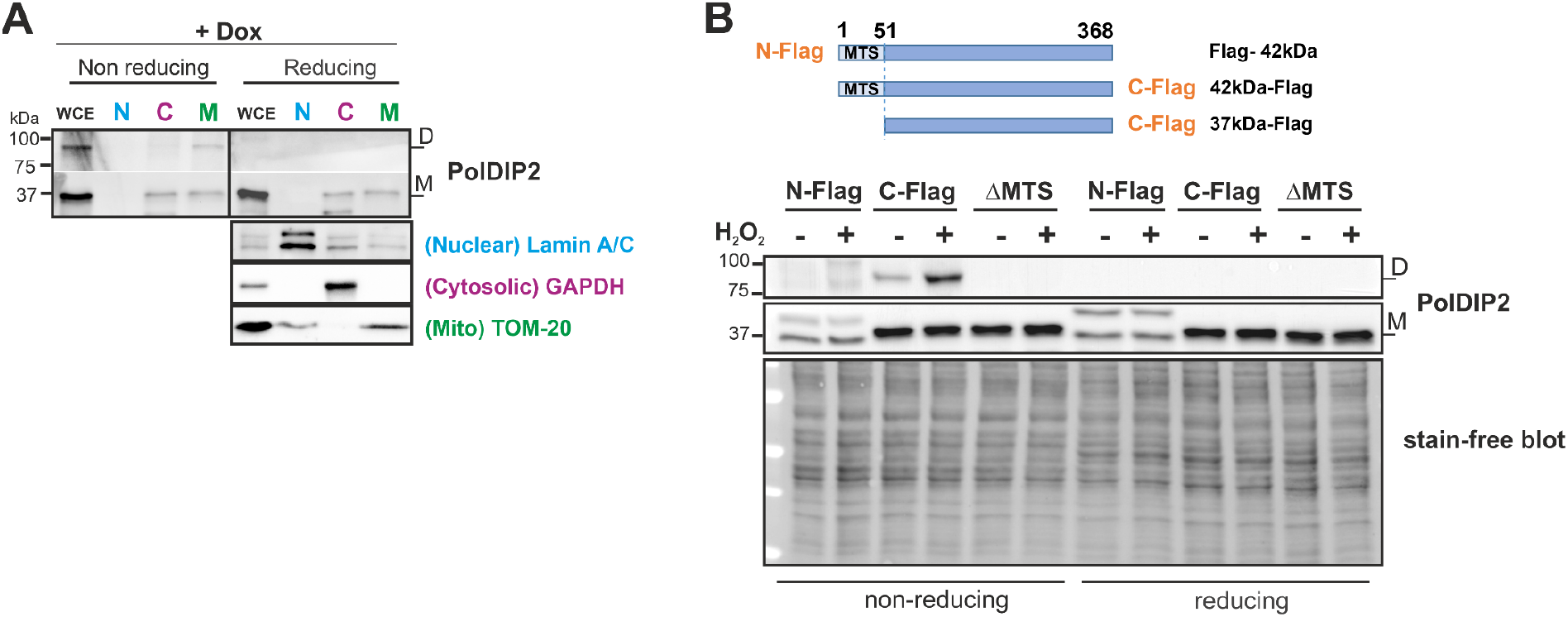
Mitochondrial localization is required for PolDIP2 dimerization. **(A).** Cell fractionation analysis shows that while monomeric PolDIP2 is present in both the cytoplasmic and mitochondrial fractions, the dimeric form is exclusively detected in mitochondria. Lamin A/C, GAPDH, and Tom20 were used as markers to confirm the purity of nuclear, cytoplasmic, and mitochondrial fractions, respectively. **(B).** Only mitochondrial-targeted and properly processed PolDIP2 forms dimers upon oxidative stress. HEK293T cells were transiently transfected with various PolDIP2 constructs: N-terminally or C-terminally Flag-tagged full-length PolDIP2, and a C-terminally Flag-tagged variant lacking the mitochondrial targeting sequence (ΔMTS). Cells were treated with 500 µM H_2_O_2_ for 20 minutes prior to harvesting. Dimerization was observed only in the C-terminally tagged, mitochondrially localized PolDIP2 following oxidative stress.

To further confirm the mitochondrial localization of PolDIP2 dimers, we transfected HEK293T cells with various PolDIP2 constructs: full-length PolDIP2 with either N-or C-terminal Flag tags, and a C-terminally Flag-tagged PolDIP2 lacking the mitochondrial targeting sequence. Placement of the Flag tag at the N-terminus prevented MTS cleavage, resulting in retention of the full-length 42-kDa precursor in the cytoplasm of the transfected cell (**Fig. S3B**), which failed to dimerize even upon H_2_O_2_ treatment (**Fig. 4B**). Similarly, the MTS-deleted PolDIP2 construct was not localized to mitochondria (**Fig. S3B**) and failed to form dimers, regardless of oxidative stress. In contrast, only the C-terminally Flag-tagged full-length PolDIP2, which was properly imported into mitochondria and processed to the 37-kDa mature form, exhibited dimer formation that was enhanced by H_2_O_2_ treatment (**Fig. 4B**). Together, these findings show that PolDIP2 dimerization occurs exclusively within mitochondria and requires correct mitochondrial import, suggesting that dimer formation is linked to mitochondrial function.

### PolDIP2 dimerization does not alter PrimPol stimulation or mtDNA replication in cells

PrimPol, a primase-polymerase, contributes to the restart of stalled replication forks and thereby supports nuclear and mitochondrial DNA integrity ^25,26^. PrimPol, is non-processive, exhibits relatively low DNA affinity and displays limited DNA synthesis activity *in vitro*^6,16,27,28^. Multiple studies, including our own, have shown that PolDIP2 enhances PrimPol’s processivity and polymerase activity^6,29^.

We therefore examined whether PolDIP2 dimerization influences PrimPol stimulation. In primer-extension assays, recombinant PolDIP2 (residues 51–368) increased PrimPol activity (**Fig S4A and S4B,** compare lane 1 with lane 2-4), and the purified dimeric protein stimulated PrimPol to a similar extent as the monomer (**Fig. S4B,** compare lanes 2-4 with lanes 6-8). Thus, Cys143-dependent dimerization does not affect PolDIP2-mediated activation of PrimPol *in vitro*.

Because PolDIP2 has also been implicated in additional mtDNA-associated processes^9,30^, we next asked whether dimerization influences mtDNA replication in cells. Using two-dimensional neutral agarose gel electrophoresis (2D-NAGE), we examined replication intermediates in HEK293 Flp-In T-REx cells expressing wild-type or C143A PolDIP2 (**Fig. S4C-E**). Doxycycline-induced overexpression of wild-type PolDIP2 caused a modest increase in the abundance of replication intermediates relative to uninduced controls (**Fig. S4E**), consistent with a mild slowing of mtDNA replication fork progression or an accumulation of replication structures. Overexpression of the dimer-deficient C143A variant produced a closely similar pattern (**Fig. S4E**). Together with our biochemical data, these results indicate that PolDIP2 overexpression can modestly influence mtDNA replication dynamics, but this effect is independent of Cys143-dependent dimerization. Thus, PolDIP2 dimerization does not make a major contribution to its roles in mtDNA-associated processes.

### PolDIP2 associates with mitochondrial proteins, including CHCHD2

Given the redox sensitivity of PolDIP2 specifically within mitochondria, we next investigated whether it associates with distinct mitochondrial protein partners that might participate in or be influenced by its redox-regulated dimerization. To define the PolDIP2 interactome in mitochondria, we performed immunoprecipitation (IP) of PolDIP2-Flag followed by mass spectrometry (MS) analysis in HEK293 cells.

Among the proteins identified, we detected previously reported mitochondrial interactors of PolDIP2, including the two components CLPX and CLPP of CLPXP protease, as well as ALAS1 (**Table S1**), validating the robustness of our IP–MS approach^12,31^. In addition to these known associations, three mitochondrial proteins not previously confirmed as PolDIP2 interactors were identified: CHCHD2, HSP60, and LONP1 (**Table S1**). These findings suggested that PolDIP2 engages in an even broader network of mitochondrial protein interactions than previously characterized.

To validate the mitochondrial protein associations identified by IP–MS, we performed co-immunoprecipitation (co-IP) of PolDIP2-Flag from doxycycline-inducible HEK293 cells and assessed selected candidates by immunoblotting. CLPX, CLPP, HSP60, LONP1, and CHCHD2 were detected in PolDIP2 immunoprecipitates (**Fig. 5A)**, confirming that PolDIP2 associates with these mitochondrial proteins in cells.

**Figure 5.**
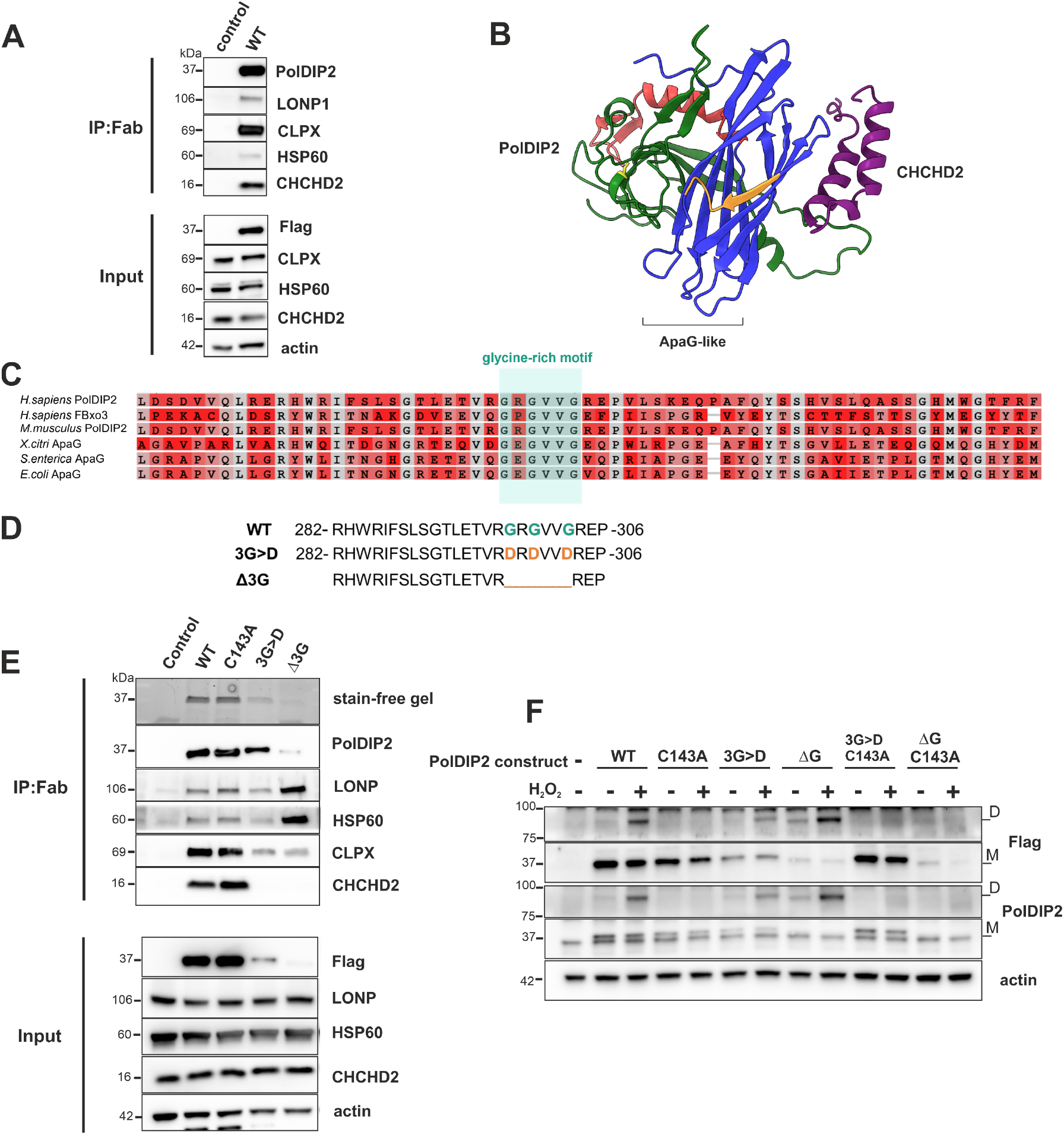
PolDIP2 interacts with several mitochondrial proteins including CHCHD2. (**A**). Western blot analysis of input and pull-down (IP:Flag) fractions from Fab-Trap immunoprecipitation in Flp-In T-Rex parental (control) or PolDI2 wild-type (WT)-expressing cells. The blots were probed with indicated antibodies. (**B**). Cartoon depiction of the Alphafold 3 predicted model for interactions between PolDIP2 (the YccV-like domain in green, ApaG-like domain in blue, Cys143 in yellow and rendered as stick) and CHCHD2 (purple), with the glycine-rich motif of PolDIP2 highlighted in orange. The unstructured, low-confidence regions of both proteins have been hidden for clarity. (**C**) Protein sequence alignment of the C-terminal domain of human PolDIP2 and other ApaG-like domain containing proteins indicates a highly conserved glycine-rich motif. The references of aligned sequence were *Homo sapiens* PolDIP2 (Q9Y2S7), *Homo sapiens* FBxo3 (Q9UK99), *Mus musculus* PolDIP2 (Q91VA6), *Xanthomonas axonopodis pv. citri* ApaG (Q8PP26), *Salmonella typhimurium* ApaG (Q56017), *Escherichia coli* ApaG (P62672). The position of the putative glycine motif (boxed in green), G^298^x^G300^xxG^303^ in human PolDIP2 is indicated. Conserved residues are shown in grey, and the shades of red colour indicate conservation levels of other residues across representative proteins. (**D**) The amino acid sequence of PolDIP2 variants harbouring mutations in the glycine-rich motif used in this study. The triple mutations 3G>D (G 298D/G300D/G303D) and deletion of the motif (ΔG298-G303) were introduced into FL-PolDIP2. (**E**) Western blot analysis of input and pull-down (IP:Flag) fractions from Fab-Trap immunoprecipitation in Flp-In T-Rex parental (control) cells or PolDIP2 cells expressing either the WT or its mutants. The blots were probed with indicated antibodies. Stain-free gel shows the enriched PolDIP2 proteins on SDS-PAGE. (**F**) Western blot analysis using anti-Flag and anti-PolDIP2 antibodies in HEK293T cells transfected with either WT or different PolDIP2 variants with or without peroxide treatment showed dimer formation under non-reducing conditions.

### A conserved glycine-rich motif in the ApaG-like domain is required for PolDIP2–CHCHD2 interaction

Among the newly validated interactors, we prioritized CHCHD2 for mechanistic follow-up, as it differs from the broadly acting mitochondrial chaperonin HSP60 or the protease LONP1 by functioning as a regulatory protein involved in redox-sensitive stress signaling. This provided a potential mechanistic link between CHCHD2 and the redox-regulated oligomerization and mitochondrial function of PolDIP2

To gain structural insight into the PolDIP2–CHCHD2 interaction detected in our proteomic screens, we generated AlphaFold3 complex predictions for the two proteins (**Fig. 5B**). Although the overall model confidence was moderate (pTM: 0.61, ipTM), in part lowered by the presence of large unstructured regions in both proteins. However, the predicted interface for the structured alpha helices of CHCHD2 consistently localized to a surface on the C-terminal ApaG/DUF525 domain of PolDIP2, adjacent to the conserved glycine-rich motif, and displayed very low expected position error in the PAE plot (**Fig. S5A**). Notably, predictions where this motif was deleted (ΔG) led to a substantial reduction in predicted complex confidence (ipTM: 0.23) (**Fig. S5B**), but not overall model confidence (pTM: 0.56), consistent with the hypothesis that this region contributes to the interaction. While the predictions do not show direct contacts between residues of the glycine-rich motif and CHCHD2, they support a model in which this motif helps shape the local surface through which PolDIP2 can engage CHCHD2.

The predicted interface lies within the C-terminal ApaG/DUF525-like domain of PolDIP2, a conserved β-sandwich fold shared with bacterial ApaG proteins and several eukaryotic F-box proteins^10,12^ (**Fig. S5**). In the PolDIP2 crystal structure^11^, this domain forms a hydrophobic groove reminiscent of the substrate-binding pocket of Fbxo3. The glycine-rich motif (GxGxxG; residues 298–303) in human PolDIP2, positioned at the base of this groove and conserved across DUF525/ApaG proteins (**Fig. 5C)** likely contributes to shaping this interaction surface. Although glycine-rich sequences in other protein families can mediate nucleotide binding^4,32^, such activity has not been observed for DUF525/ApaG homologues^11,33^, suggesting instead a role in specifying protein contacts within this domain.

To assess the role of this motif in PolDIP2 protein interactions, we generated two variants that disrupt the GxGxxG sequence: a triple glycine-to-aspartate substitution (3G>D; G298D/G300D/G303D) and a deletion of the motif (Δ3G; Δ298–303) (**Fig. 5D**). When expressed in HEK293 Flp-In T-REx cells, both mutants, particularly Δ3G, showed reduced steady-state levels compared to wild-type PolDIP2, suggesting that alteration within this region affects protein stability or expression (**Fig. 5E**). Co-IP signals for these mutants were therefore interpreted relative to input levels.

Despite their lower abundance, most tested mitochondrial interactors were recovered in co-IP at levels comparable to the wild type when normalized to enriched PolDIP2 level. In contrast, CHCHD2 binding was completely lost for both 3G>D and Δ3G mutants (**Fig. 5F**), indicating that the conserved GxGxxG motif is specifically required for PolDIP2–CHCHD2 association rather than for general mitochondrial protein interactions.

### Disruption of the glycine-rich motif enhances PolDIP2 dimerization

Given the loss of CHCHD2 binding in the glycine-rich motif mutants, we asked whether these mutations also affect PolDIP2 dimerization. Strikingly, both 3G>D and Δ3G variants displayed markedly increased levels of dimeric PolDIP2 compared with wild type, despite having substantially lower total protein abundance (**Fig. 5F**). When normalized to input levels, the relative proportion of dimer to monomer was strongly elevated for both mutants, most prominently for the Δ3G variant. H_2_O_2_ treatment further increased dimer formation in the motif mutants, indicating that oxidative conditions can still promote dimerization in their altered background.

Importantly, introduction of the C143A substitution into the 3G>D or Δ3G backgrounds abolished dimer formation, demonstrating that the enhanced dimers arise via the same Cys143-dependent intermolecular disulfide bond as in wild-type PolDIP2 (**Fig. 5F**). Together, these results indicate that the conserved glycine-rich motif not only supports PolDIP2–CHCHD2 association but also influences the redox-dependent dimerization of PolDIP2, suggesting a potential link between PolDIP2 interaction state and its oligomeric conformation.

To further test whether the oligomeric state of PolDIP2 influences its interaction with CHCHD2, we compared co-immunoprecipitation profiles of wild-type PolDIP2 with those of the dimer-deficient C143A mutant. As expected, the C143A mutation increased the monomeric pool of PolDIP2. When normalized to expression levels, recovery of most mitochondrial interactors was similar between WT and C143A. In contrast, CHCHD2 was consistently enriched in C143A co-IPs (**Fig. 5E**), indicating a preferential association with monomeric PolDIP2. These observations are in line with the model that disruption of dimerization enhances the available monomeric pool and thereby promotes CHCHD2 binding, reinforcing that CHCHD2 interacts selectively with the monomeric form of PolDIP2.

## DISCUSSION

Our study identifies redox-dependent dimerization as a previously unrecognized biochemical property of human PolDIP2 and characterizes structural determinants that regulate this process. We show that PolDIP2 forms a disulfide-linked homodimer mediated by the highly conserved Cys143, and that this oligomeric state is observed only within mitochondria. The absence of dimer formation in the cytosol and its enhancement under oxidative conditions indicate that the mitochondrial redox environment supports or promotes disulfide linkage between PolDIP2 monomers. The unique confinement of dimerization to mitochondria suggests that redox-sensitive structural changes of PolDIP2 occur only after its import, providing insight into how the protein may be regulated within the organelle.

The increase in PolDIP2 dimer formation following H_2_O_2_ (**Fig. 3A and 3B**) or rotenone exposure (**Fig. 3C**), together with the loss of this response in the Cys143A variant (**Fig. 2C and S2A**), indicates that PolDIP2 can sense changes in mitochondrial redox state through a defined cysteine-dependent mechanism.

Although Cys143-dependent dimerization did not measurably alter PolDIP2-dependent stimulation of PrimPol *in vitro* (**Fig. S4B**) or substantially change mtDNA replication dynamics in cells (**Fig. S4D**), our data instead indicate that redox-linked structural changes primarily modulate PolDIP2’s interaction profile within mitochondria rather than its potential cofactor function at replication forks. The differential behavior of monomeric and dimeric PolDIP2 in co-immunoprecipitations supports this idea, suggesting that redox conditions may help determine which interaction states of PolDIP2 are favored *in vivo* (**Fig. 5E**).

Our proteomic and biochemical analyses extend the known set of mitochondrial PolDIP2 interactors and reveal associations with HSP60, LONP1, and CHCHD2 (**Table 1**). The recovery of these proteins alongside earlier identified interactors CLPX/CLPP^12^ and ALAS1^31^ indicates that PolDIP2 is integrated into a broader interaction network than previously appreciated. The apparent enrichment of HSP60 and LONP1 in PolDIP2-ΔG co-immunoprecipitates (**Fig. 5E**) likely reflects the reduced stability of this mutant, which may alter its folding or exposure of interaction surfaces, although indirect effects cannot be excluded. We therefore interpret these associations cautiously and do not infer a direct role for PolDIP2 in mitochondrial protein quality control.

Among the newly detected interactors, CHCHD2 is particularly notable. CHCHD2 is an intermembrane space protein implicated in cristae maintenance, mitochondrial respiration, and oxidative-stress responses^13,14,34^. Pathogenic variants in CHCHD2 have also been linked to a subset of familial Parkinson’s disease cases^35^. Our interactome data are supported by two independent proteomic studies reporting enrichment of PolDIP2 in CHCHD2 pulldowns^36,37^, although these studies did not further characterize the interaction. The present work therefore provides the first mechanistic evidence defining how the two proteins associate and identifies structural features of PolDIP2 required for this interaction.

Specifically, we find that a conserved glycine-rich motif in the PolDIP2 ApaG-like domain is required for CHCHD2 binding (**Fig. 5E**), and that its disruption abolishes this interaction while markedly increasing PolDIP2 dimer formation (**Fig. 5F**). These observations suggest a direct relationship between interaction specificity and oligomeric equilibrium: CHCHD2 preferentially associates with monomeric PolDIP2, while motif-disrupting ΔG mutants accumulate disproportionately as dimers (**Fig. 5E and 5F**). AlphaFold-3 structural predictions further support this interpretation, in the wild-type model, the highest-confidence PolDIP2– CHCHD2 interface maps to the glycine-rich region (**Fig. S5A**), whereas in models lacking this motif (ΔG) the predicted interface confidence is markedly reduced (**Fig. S5B**). Although purely computational, these predictions are consistent with our biochemical evidence that CHCHD2 preferentially engages monomeric PolDIP2 via this region.

Further supporting this model, the dimer-deficient C143A variant showed increased recovery of CHCHD2 in co-immunoprecipitation assays (**Fig. 5E**). Because this mutation shifts PolDIP2 toward a monomeric state (**Fig. 2C and 2D**) without broadly altering its interaction profile, the selective enrichment of CHCHD2 is consistent with a clear preference for the monomeric form. Together with the reduced CHCHD2 binding observed for glycine-rich motif mutants, these results provide reciprocal evidence that PolDIP2’s oligomeric state directly influences its ability to engage CHCHD2.

Given CHCHD2’s role in cristae maintenance and stress signaling, and the fact that mitochondrial nucleoids frequently localize adjacent to cristae-rich regions of the inner membrane^38^, the PolDIP2–CHCHD2 interaction may take place in these cristae-adjacent microdomains. This would position PolDIP2 at sites where nucleoids, cristae architecture and local redox conditions intersect. This spatial arrangement is consistent with PolDIP2’s previously reported nucleoid associations^9^ and with our finding that its oligomeric state is redox-sensitive, raising the possibility that redox-dependent dimerization modulates which binding partners PolDIP2 engages in these regions^39^. Although we have not directly visualized PolDIP2-CHCHD2 co-localization, or demonstrated functional consequences of the interaction, our data are compatible with a model in which PolDIP2 operates at or near cristae– nucleoid interfaces, where changes in redox environment and membrane organization are tightly coordinated. Clarifying this will require targeted imaging and functional studies.

## CONCLUSIONS

This study establishes that PolDIP2 undergoes Cys143-dependent, redox-regulated dimerization exclusively in mitochondria and identifies a conserved glycine-rich motif required for its interaction with CHCHD2. While dimerization does not influence the stimulation of PrimPol, or mtDNA replication in general, the coupling between oligomeric state and interaction specificity suggests that PolDIP2 may use redox-sensitive structural transitions as part of its regulatory behavior within mitochondria. These findings refine our understanding of PolDIP2’s biochemical properties and provide a framework for future work aimed at defining its roles in mitochondrial homeostasis.

## DATA AVAILABILITY

The structure has been deposited in the Protein Data Bank with PDB Code: XXXX

## SUPPLEMENTARY DATA ACKNOWLEDGMENTS

We acknowledge Annika Thorsell (Proteomics Core Facility at Sahlgrenska Academy, Gothenburg University Sweden) for providing assistance with Mass spectrometry. We thank Mikael Lindberg and the Protein Expertise Platform (PEP-Umeå University) for PolDIP2 WT plasmid cloning. We also thank Noopur Singh and Josefin Forslund for the helpful discussion.

## Supplementary Data

**Table S1.**
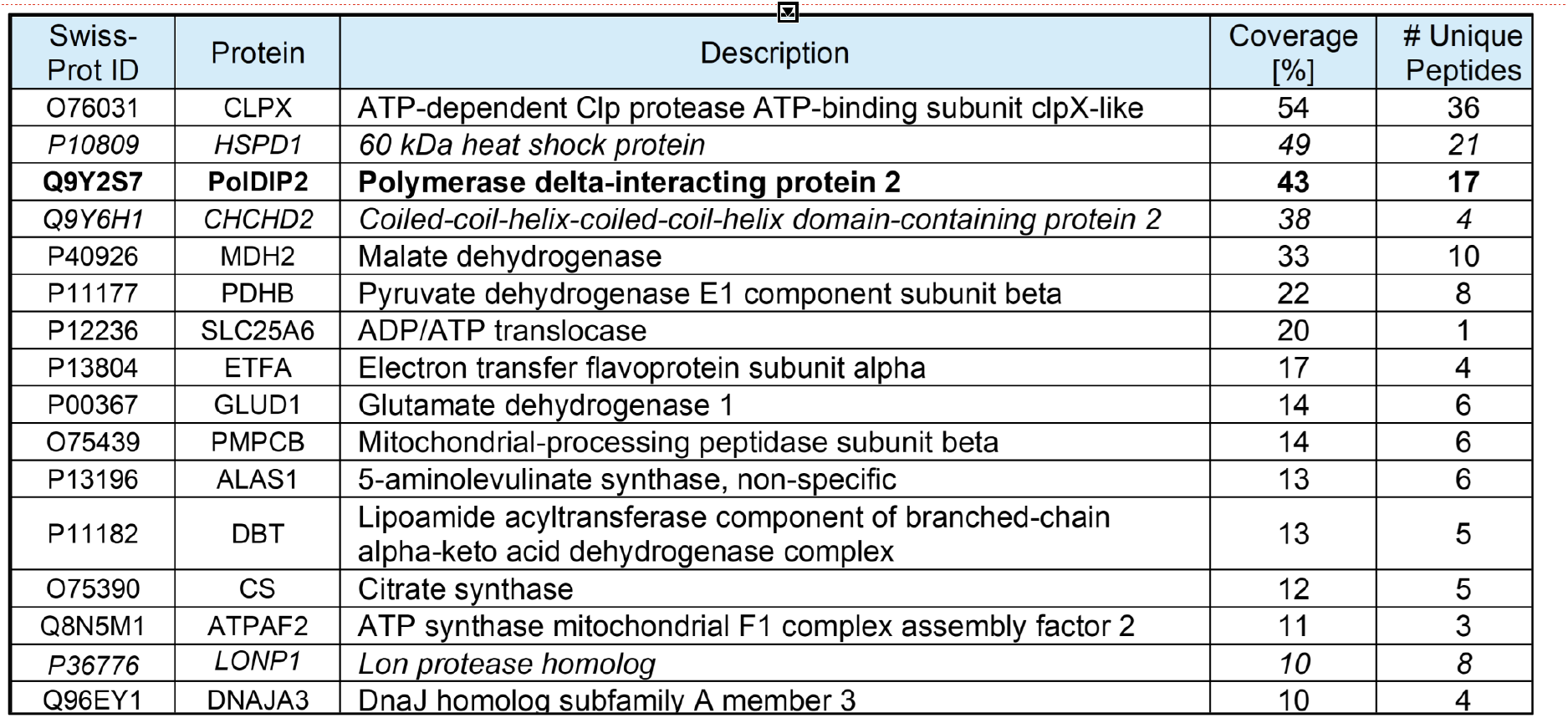
Proteins co-purified with WT PolDIP2-Flag in co-immunoprecipitation.

| Swiss-Prot ID | Protein | Description | Coverage [%] | # Unique Peptides |
| --- | --- | --- | --- | --- |
| O76031 | CLPX | ATP-dependent Clp protease ATP-binding subunit clpX-like | 54 | 36 |
| P10809 | HSPD1 | 60 kDa heat shock protein | 49 | 21 |
| **Q9Y2S7** | **PolDIP2** | **Polymerase delta-interacting protein 2** | **43** | **17** |
| Q9Y6H1 | CHCHD2 | Coiled-coil-helix-coiled-coil-helix domain-containing protein 2 | 38 | 4 |
| P40926 | MDH2 | Malate dehydrogenase | 33 | 10 |
| P11177 | PDHB | Pyruvate dehydrogenase E1 component subunit beta | 22 | 8 |
| P12236 | SLC25A6 | ADP/ATP translocase | 20 | 1 |
| P13804 | ETFA | Electron transfer flavoprotein subunit alpha | 17 | 4 |
| P00367 | GLUD1 | Glutamate dehydrogenase 1 | 14 | 6 |
| O75439 | PMPCB | Mitochondrial-processing peptidase subunit beta | 14 | 6 |
| P13196 | ALAS1 | 5-aminolevulinate synthase, non-specific | 13 | 6 |
| P11182 | DBT | Lipoamide acyltransferase component of branched-chain alpha-keto acid dehydrogenase complex | 13 | 5 |
| O75390 | CS | Citrate synthase | 12 | 5 |
| Q8N5M1 | ATPAF2 | ATP synthase mitochondrial F1 complex assembly factor 2 | 11 | 3 |
| P36776 | LONP1 | Lon protease homolog | 10 | 8 |
| Q96EY1 | DNAJA3 | DnaJ homolog subfamily A member 3 | 10 | 4 |

**Table S2.** Crystallography table for PolDIP2 dimer.

| **Data collection summary** | **PolDIP2** |
| --- | --- |
| Resolution range | 56.01 – 2.6 (2.77-2.61) |
| Space group | P 2<sub>1</sub> 2<sub>1</sub> 2<sub>1</sub> |
| Cell dimensions |  |
| a, b, c (Å) | 70.49 92.23 93.35 |
| $\alpha$, $\beta$, $\gamma$ (°) | 90 90 90 |
| Total reflections | 250200 (38373) |
| Unique reflections | 19085 (2933) |
| Multiplicity | 13.1(13.0) |
| Completeness (%) | 99.4 (97.2) |
| Mean I/sigma (I) | 21.4 (3.66) |
| R-means | 0.165 (0.985) |
| CC(1/2) | 0.998 (0.910) |
| **Refinement summary** |  |
| Resolution range | 56.01-2.8 (2.98-2.8) |
| R-work | 0.274 (0.332) |
| R-free | 0.304 (0.379) |
| Number of non-hydrogen atoms |  |
| protein | 4154 |
| solvent | 29 |
| RMS (bonds) | 0.002 |
| RMS (angles) | 0.67 |
| Ramachandran favored (%) | 93.8 |
| Ramachandran allowed (%) | 6.2 |
| Ramachandran outliers (%) | 0.00 |
| Average B-factor | 63.6 |
| protein | 64.29 |
| solvent | 46.0 |

## Supplementary Figures

**Supplementary Figure 1.**
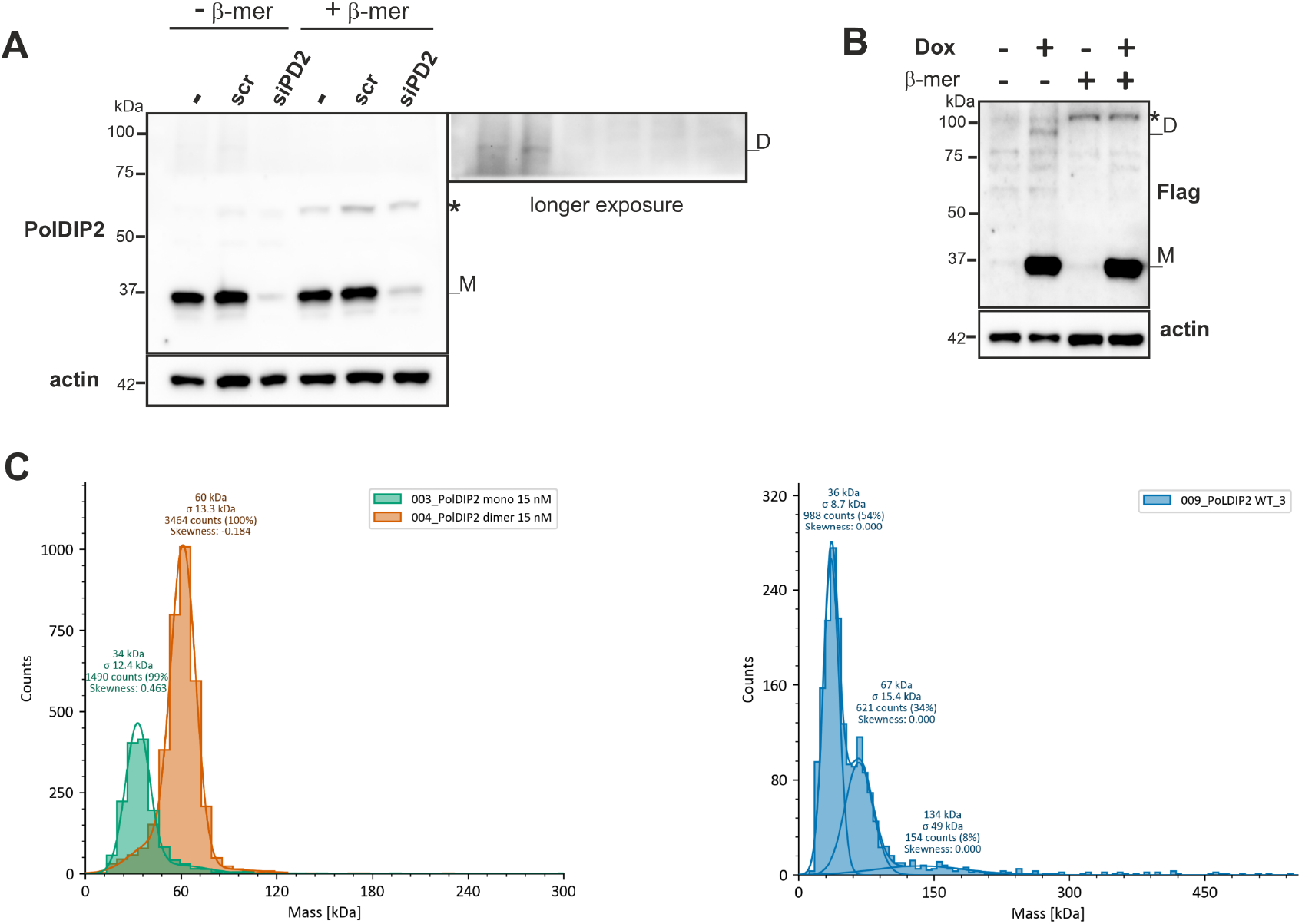
**(A)** Western Blot analysis of HEK293T cells transfected with siRNA against PolDIP2 or scramble siRNA for 72 hours. siRNA silencing of PolDIP2 effectively depleted endogenous PolDIP2 monomers. The less abundant oligomer was absent in the presence of β-mercaptoethanol and similarly reduced upon RNAi knockdown. Both short (left) and long (right) exposures of the blot are shown. **(B)** Western blot analysis using an anti-Flag antibody detected C-terminally Flag-tagged PolDIP2 expressed in Flp-In™ 293 T-REx cells, with and without doxycycline induction, under reducing and non-reducing conditions. *non-specific band. **(C)** Mass photometry (MP) analysis showed that the truncated recombinant human PolDIP2 monomer and dimer gave two distinct profiles of mass distribution, 34 ± 12.4 kDa and 60 ± 13.3 kDa respectively (left panel). Overexpressed WT PolDIP2-Flag enriched from cultured cells with Flag-affinity beads was also run in MP. Two populations separated by size difference were detected: the major species with a mass of 36 ± 8.7 kDa corresponds to the monomer while the minor species (67 ± 15.4 kDa) corresponds to the less abundant dimers (right panel).

**Supplementary Figure 2.**
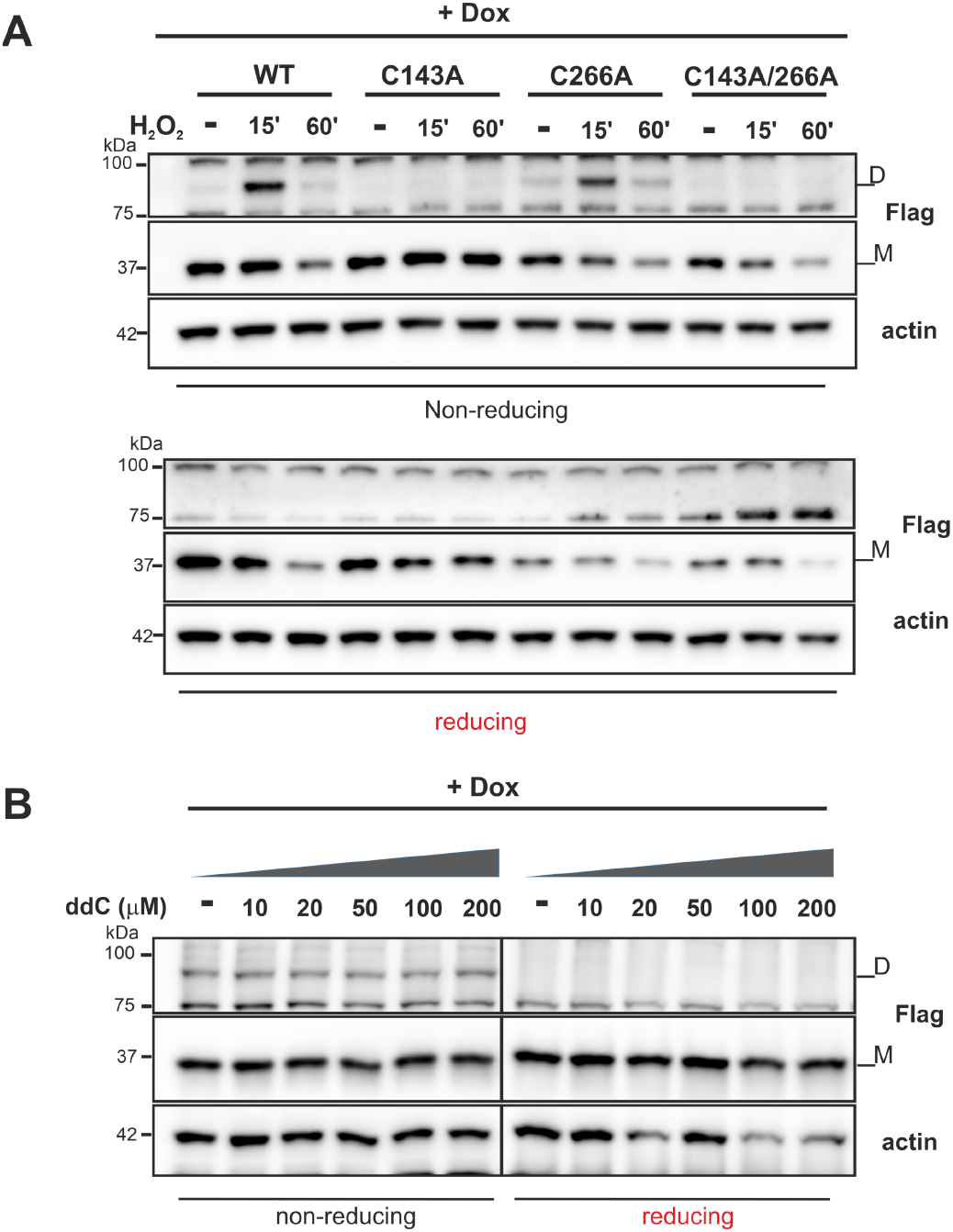
**(A)** Substitution of Cysteine 266 to Alanine did not disrupt PolDIP2 dimer formation in cells. Western blot analysis using anti-Flag antibody in HEK293T cells transfected with wild-type (WT) and different PolDIP2 variants with or without peroxide treatment showed dimer formation under non-reducing conditions. PolDIP2 dimer formation was abolished when mutating Cysteine 143 while substitution of Cysteine 266 to Alanine did not appear to affect protein dimerization. (**B**) Western blot analysis using anti-Flag antibody in Flp-In™ 293 T-REx cells expressing WT-PolDIP2 after treating with increasing concentrations of 2’,3’-dideoxycytidine (ddC) for 24 hours. It was shown that ddC exposure did not enhance PolDIP2 dimer formation

**Supplementary Figure 3.**
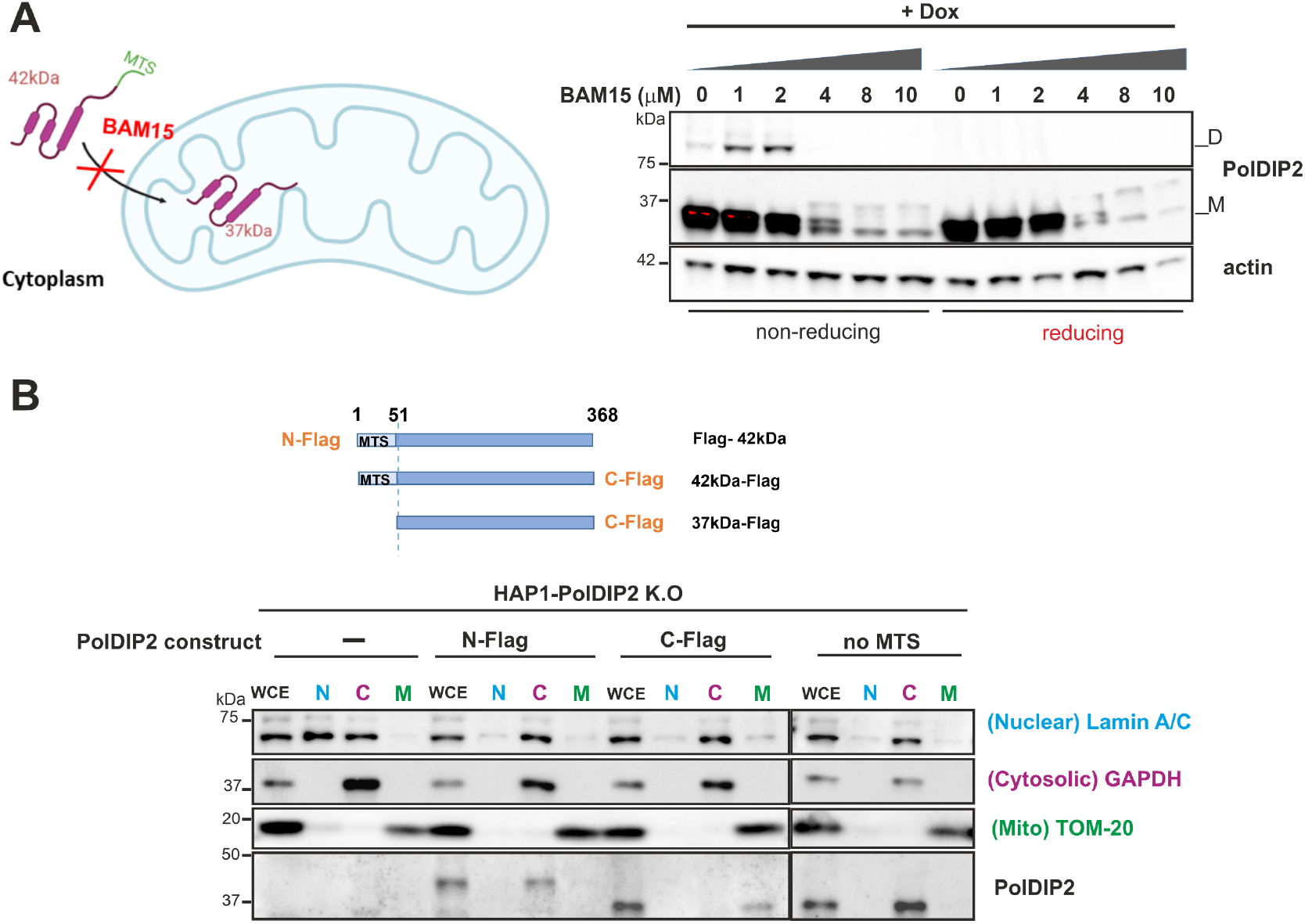
**(A)** Schematic illustration of simplified action of the mitochondrial uncoupler BAM15 on PolDIP2 localization (left). BAM15 which disrupts mitochondrial membrane potential (MPP) that drives protein import into mitochondria. Western blot analysis (right) in Flp-In™ 293 T-REx cells expressing WT-PolDIP2 treated with increasing concentrations of BAM15 for 18 hours. The mitochondrial PolDIP2 variant of 37kDa disappeared and the premature PolDIP2 (42kDa) was detected upon BAM15 treatment, indicating that PolDIP2 import into mitochondria is MPP-driven. Moreover, the dimeric species was similarly lost upon treatment, providing more evidence of mitochondrial localization of the PolDIP2 dimer. (**B**) Western blot analysis of cell fractionation showed different subcellular localizations of different PolDIP2-Flag constructs. PolDIP2-knockout HAP1 cells were transiently transfected with various PolDIP2 constructs: N-terminally or C-terminally Flag-tagged full-length PolDIP2, and a C-terminally Flag-tagged variant lacking the mitochondrial targeting sequence (ΔMTS). Flag tag at the N-terminus prevents the mitochondrial targeting and processing of PolDIP2, resulting in the retained pre-mature protein in the cytoplasm. In contrast, C-terminally Flag-tagged variant was properly imported into mitochondria and processed into the mature 37-kDa protein, whereas MTS-lacking variant failed to be imported in mitochondria but resides in the cytosol instead. Lamin A/C, GAPDH, and Tom20 were used as markers to verify the separations of nuclear, cytoplasmic, and mitochondrial fractions, respectively.

**Supplementary Figure 4.**
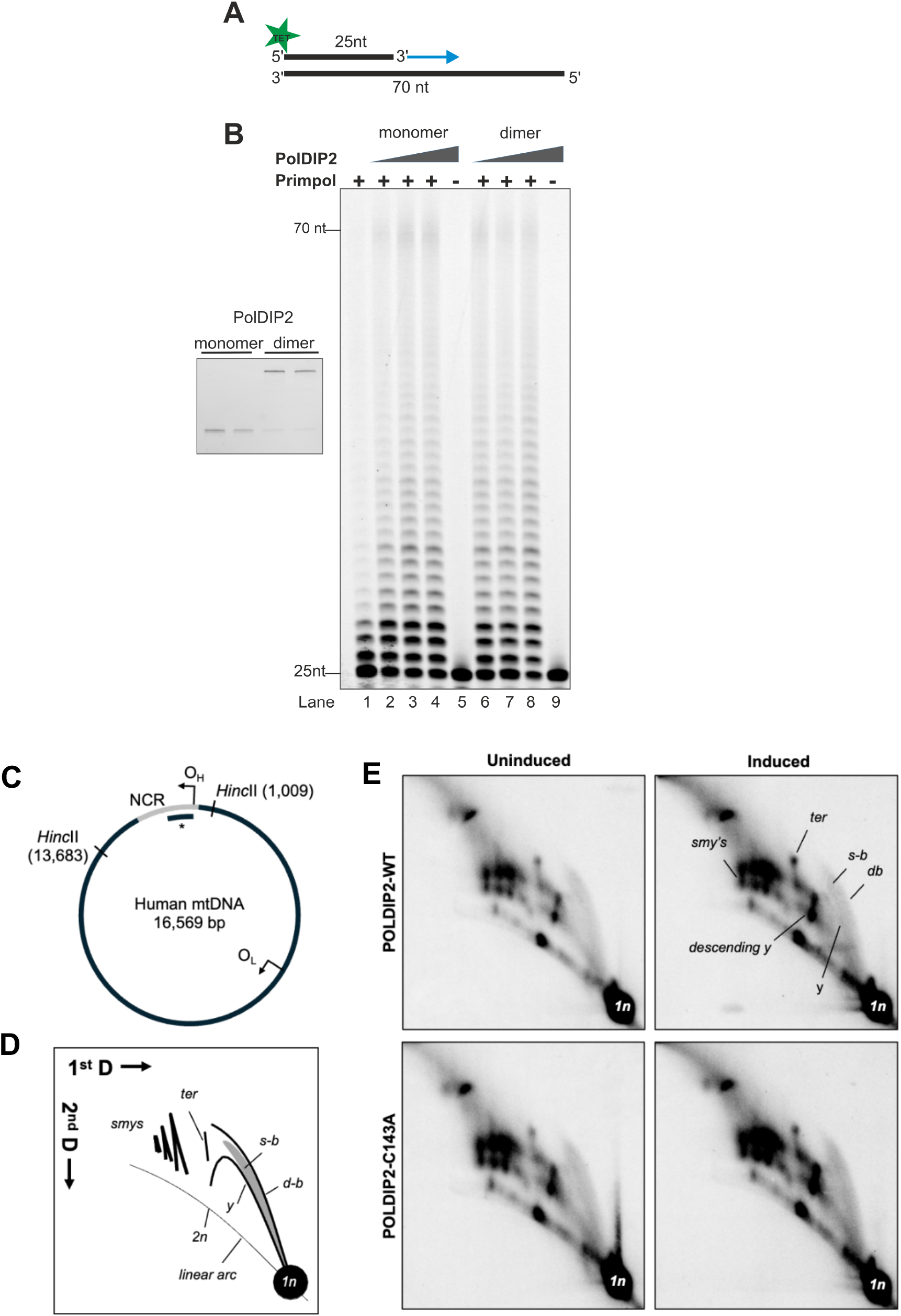
**(A)** Schematic presentation of 5’-TET labelled 25/70 nt primer/template used for primer extension assay in B. **(B)** Primer extension assay with Primpol and increasing concentrations of PolDIP2 on DNA substrate shown in A. The small panel on the right side shows purified PolDIP2 on SDS-PAGE gel stained with Instant Blue. The SDS-PAGE gel on the right shows the oligomeric forms of recombinant PolDIP2 used for the assay. PolDIP2 dimer can stimulate Primpol to a similar extent as the monomeric protein under set conditions. (**C**) A schematic representation of *Hinc*II restriction sites on mtDNA and the location of the probe (asterisk) used in 2D-AGE analysis. (**D**) Interpretation of the 2D-AGE pattern. Majority of the molecules are non-replicating double-stranded mtDNA fragments and form a 1n spot (3.9 kb *Hinc*II). Progression of the replication fork through the fragment gives rise to y-arc. Partially single-stranded bubble arc (s-b) contains molecules that are partially covered my RNA according to the strand-displacement mode of mtDNA replication. As the restriction enzyme is unable to cut RNA:DNA fragments these molecules are further transformed into y-like molecules that give rise to slow moving y-like arcs (smys) often containing traces of RNA molecules (a cloud of r-smys). Fully double-stranded bubbles (d-b) indicate an initiation of replication. Replication terminates at O_H_ within the fragment producing Holliday junctions-like molecules (ter). (**E**) Overexpression of wild type and C143A mutant POLDIP2 leads to an accumulation of all obseved replication intermediates reflective of mtDNA replication stress. In particular, the accumulation of intermediates in the descending portion of the y-arc indicates a mild replication stalling at the O_H_.

**Supplementary Figure 5.**
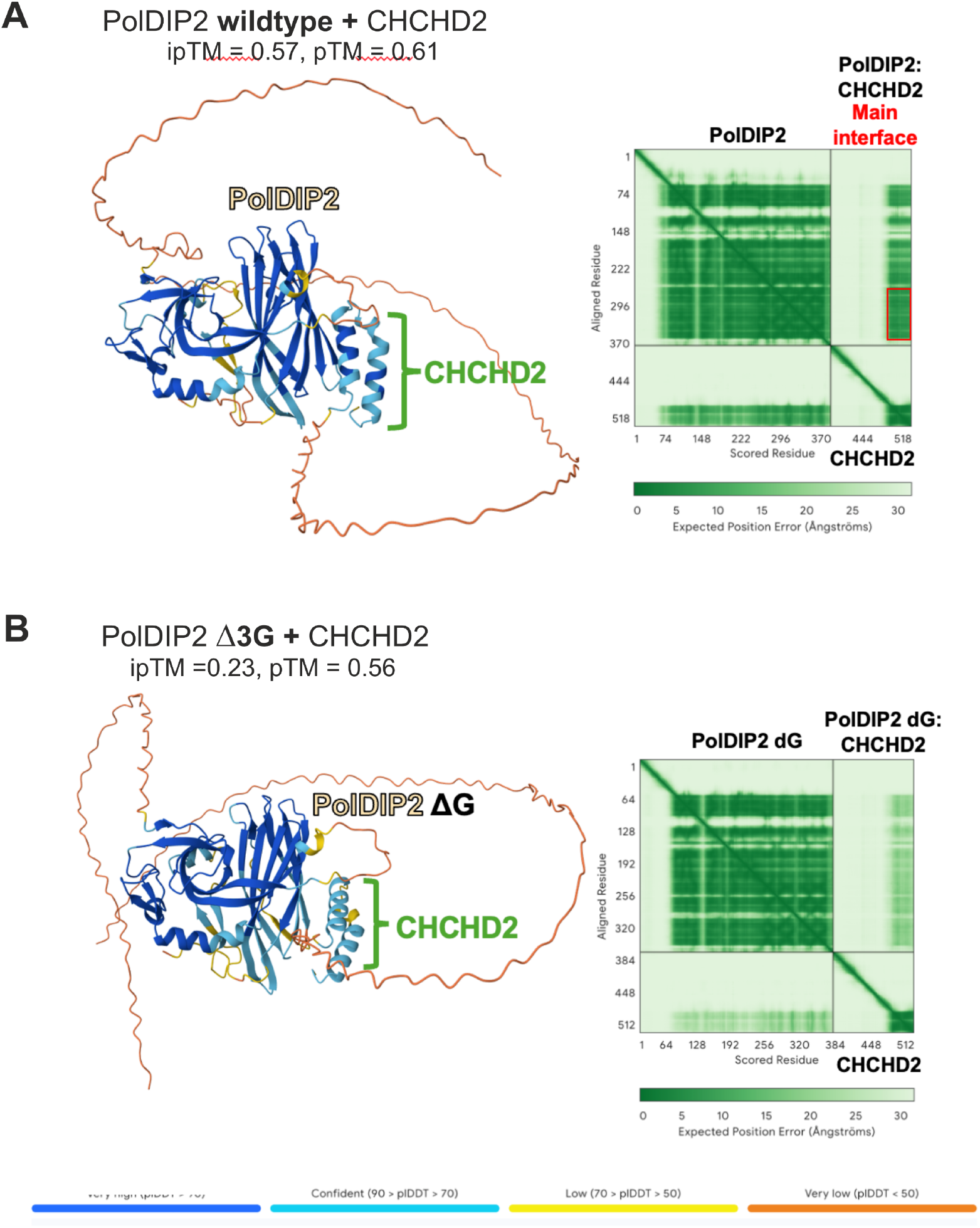
Cartoon representations of the Alphafold3 predictions for PolDIP2 (wildtype)-CHCHD2 **(A)** and PolDIP2 (deltaG)-CHCHD2 **(B)**. The cartoons are coloured by predicted local distance difference test (pLDDT), as per the key (bottom). Multimer predicted aligned error (PAE) plots are provided alongside each prediction and the region for the main interface between the two proteins is highlighted in a red box. The cartoon representations and PAE plots are direct output from the AlphaFold3 Server. Predictions were run with a random seed.

